# DISTINCTIVE FEATURES OF THE RESPIRATORY SYNCYTIAL VIRUS PRIMING LOOP COMPARED TO OTHER NON-SEGMENTED NEGATIVE STRAND RNA VIRUSES

**DOI:** 10.1101/2022.03.18.484858

**Authors:** Tessa N. Cressey, Afzaal M. Shareef, Victoria A. Kleiner, Sarah L. Noton, Patrick O. Byrne, Jason S. McLellan, Elke Muhlberger, Rachel Fearns

## Abstract

*De novo* initiation by viral RNA-dependent RNA polymerases often requires a polymerase priming residue, located within a priming loop, to stabilize the initiating NTPs. Polymerase structures from three different non-segmented negative strand RNA virus (nsNSV) families revealed putative priming loops in different conformations, and an aromatic priming residue has been identified in the rhabdovirus polymerase. In a previous study of the respiratory syncytial virus (RSV) polymerase, we found that Tyr1276, the L protein aromatic amino acid residue that most closely aligns with the rhabdovirus priming residue, is not required for RNA synthesis but two nearby residues, Pro1261 and Trp1262, were required. In this study, we examined the roles of Pro1261 and Trp1262 in RNA synthesis initiation. Biochemical studies showed that substitution of Pro1261 inhibited RNA synthesis initiation without inhibiting back-priming, indicating a defect in initiation. Biochemical and minigenome experiments showed that the initiation defect incurred by a P1261A substitution could be rescued by factors that would be expected to increase the stability of the initiation complex, specifically increased NTP concentration, manganese, and a more efficient promoter sequence. These findings indicate that Pro1261 of the RSV L protein plays a role in initiation, most likely in stabilizing the initiation complex. However, we found that substitution of the corresponding proline residue in a filovirus polymerase had no effect on RNA synthesis initiation or elongation. These results indicate that despite similarities between the nsNSV polymerases, there are differences in the features required for RNA synthesis initiation.

**Author Summary:** RSV has a significant impact on human health. It is the major cause of respiratory disease in infants and exerts a significant toll on the elderly and immunocompromised. RSV is a member of the *Mononegavirales*, the non-segmented, negative strand RNA viruses (nsNSVs). Like other viruses in this order, RSV encodes an RNA dependent RNA polymerase, which is responsible for transcribing and replicating the viral genome. Due to its essential role during the viral replication cycle, the polymerase is a promising candidate target for antiviral inhibitors and so a greater understanding of the mechanistic basis of its activities could aid antiviral drug development. In this study, we identified an amino acid residue within the RSV polymerase that appears to stabilize the RNA synthesis initiation complex and showed that it plays a role in both transcription and RNA replication. However, the corresponding residue in a different nsNSV polymerase does not appear to play a similar role. This work reveals a key feature of the RSV polymerase but identifies differences with the polymerases of other related viruses.

## Introduction

The non-segmented negative strand RNA viruses (nsNSVs) are a large and diverse order of viruses, currently comprising 11 virus families. These families include, amongst others, the filoviruses, such as Ebola and Marburg virus, the paramyxoviruses, such as parainfluenza viruses types 3 and 5 (PIV-3 and PIV-5), the rhabdoviruses, rabies and vesicular stomatitis virus (VSV) and the pneumoviruses, respiratory syncytial virus (RSV) and human metapneumovirus (HMPV) (1). The nsNSVs encode a multifunctional RNA dependent RNA polymerase that can generate capped and polyadenylated mRNAs and perform genome replication (2, 3). The polymerase is an attractive target for antiviral drug development due to its essential role in viral transcription and genome replication, coupled with the fact that facets of its activity are distinct from those of cellular enzymes (2, 4). In addition, the enzymatic active sites of nsNSV polymerases are well conserved throughout the order, raising the possibility of generating antiviral compounds with broad spectrum activity. However, despite a significant degree of primary amino acid sequence conservation within active sites, differences have been identified between nsNSV polymerases (5). Therefore, it is valuable to characterize key events in transcription and replication, as a means of understanding structure-function relationships of the polymerases and similarities and differences between viruses.

In the case of RSV, the polymerase consists of a complex of the large polymerase subunit (L), which contains the enzymatic domains for RNA synthesis, capping and cap methylation, and a cofactor, phosphoprotein (P). The L-P complex initiates both transcription and replication from a short extragenic region at the 3′ end of the genome, referred to as the *leader* (*le*) promoter region. The key promoter residues are contained within the first 12 nt of the 44-nt *le* region (6–8). This promoter contains two initiation sites, at positions 1U and 3C (9). Transcription is initiated at position 3C. In this case, the polymerase synthesizes a short (∼25 nt) le+ transcript, complementary to the *le* promoter region, and then reinitiates RNA synthesis at the *gene start* (*gs*) signal at the beginning of the first gene (5, 9). Having initiated at the *gs* signal, the polymerase caps the RNA, likely using a polyribonucleotidyltransferase activity conferred by a conserved capping domain and methylates the RNA at the 2′O and guanine N7 positions (10). Capping of the mRNA allows the polymerase to become processive and to synthesize a full-length mRNA (5, 11). Replication is initiated at position 1U. In this case, the RNA can become encapsidated as it is being synthesized (9), enabling the polymerase to elongate the RNA to the end of the genome to produce antigenome RNA. The antigenome contains a promoter at its 3′ terminus, referred to as the *trailer* (*tr*) promoter. The *tr* promoter has strong sequence similarity to the *le* promoter and likewise signals initiation from positions 1U and 3C (12, 13). In this case, initiation at 3C yields only the short ∼25 nt transcript, but no mRNAs, and initiation at 1U yields encapsidated genome RNA (14).

RSV transcription and replication are both initiated by a *de novo*, or primer independent, initiation mechanism, in which the polymerase polymerizes two NTPs to form the dinucleotide that begins the RNA chain (9, 14). Studies with other polymerases have shown that *de novo* initiation requires appropriate positioning of a number of elements within the polymerase: the RNA template, which must be positioned such that the 3′ terminus is positioned appropriately relative to the active site, two metal cations, typically Mg^2+^, and the two initiating NTPs (NTP_1_ and NTP_2_), which must enter the substrate pocket and be positioned to allow phosphodiester bond formation (15, 16). A key feature of many polymerases is a priming residue, typically an aromatic or ring-containing amino acid, which can form base-stacking interactions with the initiating NTPs (15, 17–23). This stabilizes the NTPs, facilitating *de novo* initiation. Priming residues are likely to be particularly important for initiation at the 3′ terminus of a template, compared to an internal site, due to different energy requirements for terminal versus internal initiation (22). The priming loop can also buttress the 3′ end of the template to ensure accurate initiation at the terminus and to prevent the 3′ end of the template being extended by a back-priming mechanism (18, 24). Structures are available for polymerases from three nsNSV families, the rhabdoviruses, paramyxoviruses and pneumoviruses (10, 25–31). The rhabdovirus L proteins have a loop that extends from the capping domain into the enzymatic cavity of the polymerization domain that is well positioned to be a priming loop (27–29, 32). Studies with rabies virus showed that a tryptophan residue (rabies virus L Trp1180), positioned at the tip of this loop, is required for initiation at the 3′ terminus of the template, but not at an internal initiation site located at position 7 of the promoter, indicating that this is a priming residue (20). The positioning of this loop and priming residue suggests that the rhabdovirus polymerase structures represent a pre-initiation state, in which the polymerase is poised to accept incoming RNA template and NTPs. The paramyxovirus and pneumovirus polymerases have a loop structure in the position corresponding to the rhabdovirus priming loop region in a linear sequence alignment. However, in the published structures for paramyxoviruses and pneumoviruses, the priming loop is folded out of the way of the polymerization active site and integrated with the capping domain (10, 26, 30)(Fig 1A).

**Fig 1.**
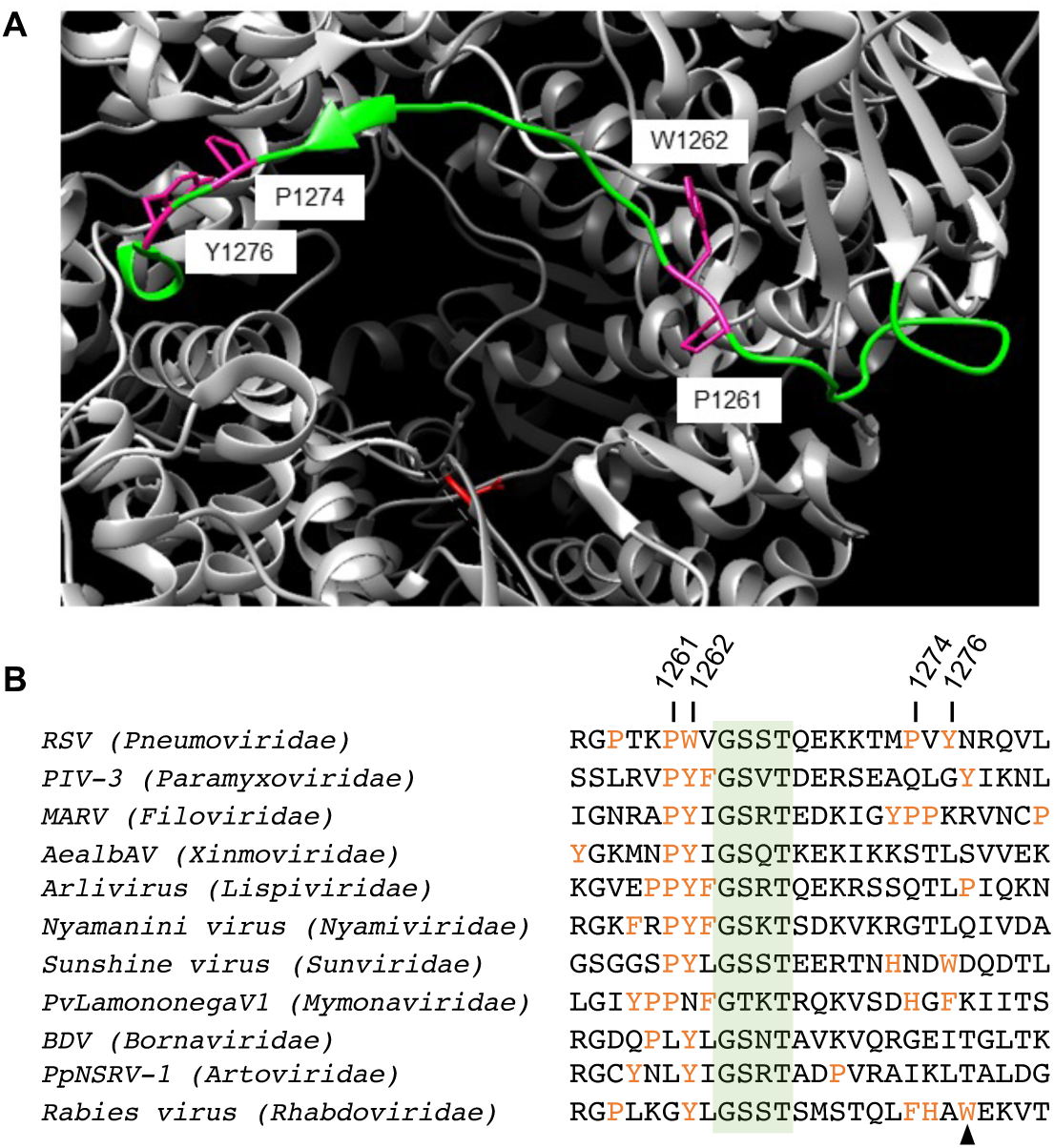
Amino acids with potential as priming residues in the nsNSVs. (A) Image illustrating the putative priming loop of the RSV L protein indicating the positions of the residues under investigation. The extended loop region is shown in green, the residues that were analyzed are in magenta, and one of the catalytic aspartic acid residues in the RNA synthesis active site is in red. The image was created using Chimera (51) using PDB: 6PZK (10). (B) Alignment of the putative priming loop region of representative virus L proteins from the eleven different families within the nsNSVs. Residues capable of forming base-stacking interactions are in orange. A conserved GxxT motif is highlighted in green. The RSV residues investigated in this study are labeled, and the tryptophan identified as the rabies virus polymerase priming residue is indicated with an arrowhead. The alignment is based on the following sequences: RSV (KT992094.1); parainfluenza virus type 3, PIV-3 (FJ455842); Lake Victoria marburgvirus, MARV (DQ447658); Aedes albopictus anphevirus, AealbAV (MW147277.1); Hubei odonate virus 10 arlivirus (NC_032944.1); Nyamanini virus (NC012703.1); Sunshine virus (NC_025345.1); plasmopara viticola lesion associated mymonavirus 1, PvLamononegaV1 (MN557002.1); Borna disease virus, BDV MT33064.1; Pteromalus puparum negative-strand RNA virus 1, PpNSRV-1 (NC_038269.1); rabies virus (EU877068).

The priming loop sequence is poorly conserved between nsNSV polymerases and there is no clear conservation of the rabies virus tryptophan priming residue (Fig 1B). According to primary sequence alignments, the residue corresponding to the rabies virus priming tryptophan in RSV L protein is Tyr1276 (Fig 1B). In a previous study, employing a cell-based minigenome assay, we mutated this and other ring-based amino acids that lie within the putative priming loop structure in the RSV polymerase. We found that mutant polymerases containing an alanine substitution at Tyr1276, or its near neighbor Pro1274, yielded RNA products at wild-type (wt) levels (30), but substitution of two nearby residues Pro1261 and Trp1262 inhibited RNA synthesis from the 1U and 3C sites, suggesting that one of these amino acid residues might represent the priming residue (30). In this study, we examined the roles of Pro1261 and Trp1262 of RSV L protein in more detail. The data obtained show that substitution of Pro1261 or Trp1262 yielded defects in initiation that could be rescued under conditions that would be expected to increase the stability of initiation complexes, indicating that these residues have a role in RNA synthesis initiation. However, substitution of the corresponding proline and aromatic residues in a filovirus polymerase had no impact on RNA synthesis initiation. Together, these findings reveal a key role for Pro1261 and Trp1262 in RNA synthesis initiation by the RSV polymerase but highlight differences between nsNSV polymerases in the structural features that are required for RNA synthesis initiation.

## Results

### The P1261A and W1262A mutants behave differently on le versus tr promoters

The early stages of RSV RNA synthesis in cells can be analyzed using a minigenome system. Specifically, by performing primer extension analysis using primers that hybridize close to the 5′ end of the RNA synthesis products, 1U and 3C initiated RNAs that are elongated a short distance within the promoter region can be analyzed. By using a minigenome limited to the first step of RNA replication, the levels of 3C initiation are unaffected by the levels of 1U (replication) initiation (9), allowing them to be analyzed as independent events. Previously, we used this approach to examine RNA synthesis by the P1261A, W1262A, P1274A and Y1276A L mutants using a minigenome containing a 3′ *tr* promoter (30). The RSV *le* and *tr* regions are almost identical in the essential promoter regions at their 3′ termini but differ at positions 4 and 12 (7, 12) (Fig 2A). Here, we also examined these L mutants on a minigenome containing a 3′ *le* promoter (Figs 2A-C and S2), with analysis of activity on the *tr* promoter shown for comparison (Fig 2D-F). The levels of mutant polymerase expression are shown in Fig S1. The mutant polymerases showed broadly similar activities on the two promoters, with the P1261A and W1262A mutants showing a more profound RNA synthesis defect than the P1274A and Y1276A mutants, as described previously (30). However, there were some promoter-specific differences in L mutant activities. Whereas the P1274A mutant was fully active on both promoters, the Y1276A mutant generated slightly less RNA from the *le* promoter than the *tr* promoter, with both 1U and 3C products being reduced to a similar extent. The P1261A and W1262A mutants also behaved differently on the two promoters, but with differential effects on RNA synthesis from position 1U versus position 3C. P1261A and W1262A did not generate RNA from position 1U of either promoter (Fig 2B, 2C, 2E, 2F). However, whereas there was barely detectable RNA synthesis from position 3C of the *tr* promoter, both mutants generated RNA at ∼25% of wt levels from position 3C of the *le* promoter (Fig 2B, 2C, 2E, 2F). Thus, changing the identity of the promoter partially restored the P1261A and W1262A 3C initiation defect.

**Fig 2.**
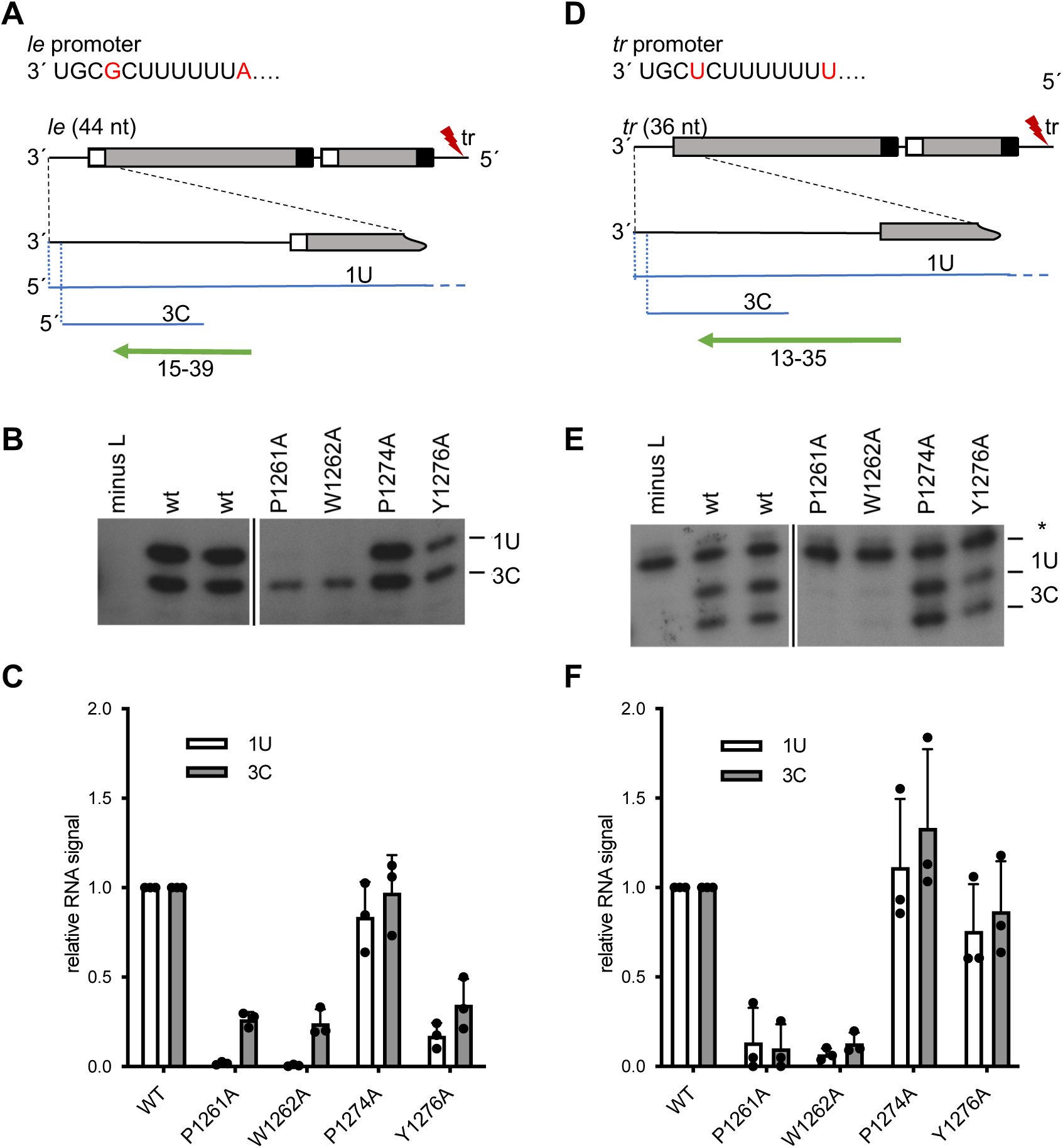
The P1261A and W1262A mutants behave differently on le versus tr promoters. (A and D) Schematic diagrams showing the organization of the minigenomes, their RNA synthesis products and the primers used for primer extension analysis. The minigenomes contained at their 3′ end either the 44-nt *le* promoter region and the first *gs* signal (A) or the 36-nt tr promoter region with no *gs* signal (D). Both minigenomes contained a mutation (indicated with a red lightning bolt) in the 5′ tr region to inactivate the *tr* promoter at the 3′ end of the replication product. An enlarged view of the 3′ end of the minigenome is shown below, the RNAs generated from the 1U and 3C sites are indicated in blue, and the primers used to detect these products are shown with green arrows. (B and E) Analysis of RNA products generated from the 1U and 3C sites in a minigenome containing either a *le* (B) or *tr* (E) promoter by mutant L polymerases. The core promoter sequences are shown, with the nucleotide differences between the *le* and *tr* promoters in red font. The RNA products were detected by primer extension analysis using the primers shown in panels A and D. The sizes of products were determined by markers shown in Fig 2S. Each panel is from the same gel, but with intervening lanes removed, as indicated by the cut. The asterisk in panel E indicates a non-specific background band. (C and F) Quantification of the data presented in panels B and E, respectively. The bars show the mean and standard deviation for three independent experiments (the data points for each experiment are shown).

### The RSV L P1261A mutant is defective in RNA synthesis from the promoter, but is capable of elongating RNA by back-priming

Although the minigenome assay used for Fig 2 can provide information regarding the ability of the polymerase to initiate at the position 1 and 3 sites, this assay does not distinguish between a global effect on RNA synthesis versus a specific defect in initiation. Therefore, to determine if either the P1261A or W1262A substitutions specifically inhibit RNA synthesis from the promoter, we utilized an *in vitro* RNA synthesis assay in which RNA synthesis activities are reconstituted using purified recombinant RSV polymerase and an RNA oligonucleotide template. It has previously been shown that if provided a template containing nucleotides 1-25 of the *tr* promoter region, RSV L-P complexes initiate RNA synthesis from the 1U and 3C sites, with position 3 products being dominant. In addition, the *tr* template can fold into a secondary structure and the RSV polymerase can extend the 3′ end of the template by one to three nucleotides, an activity referred to as back-priming (Fig 3A) (14). It is anticipated that a priming residue would be necessary for *de novo* initiation at 1U and possibly 3C but would not be required for back-priming elongation activity. Thus, this assay allows us to distinguish between effects on initiation versus elongation within the same reaction.

**Fig 3.**
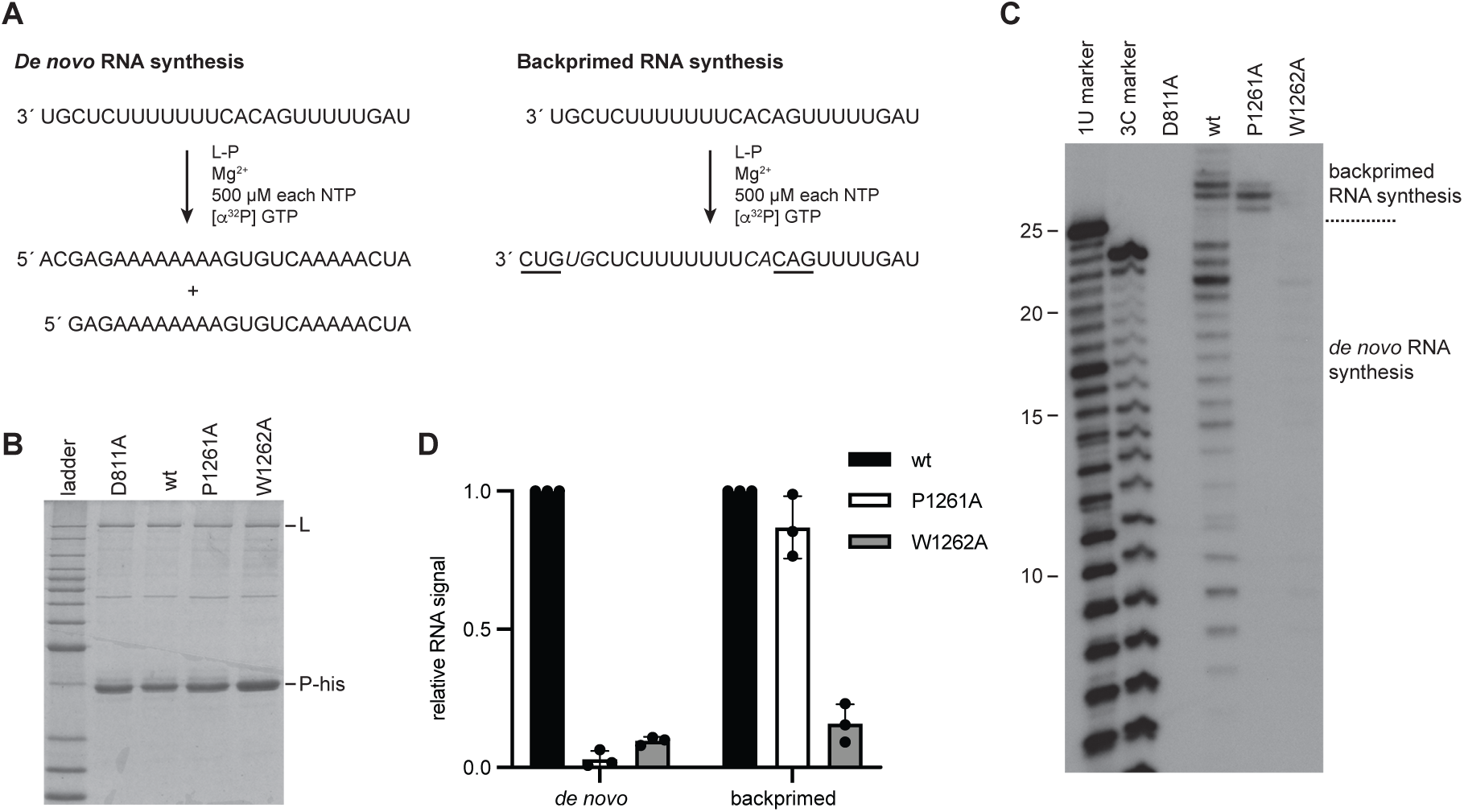
Substitution of P1261 inhibits RNA synthesis from the promoter, but not RNA elongation by back-priming. (A) Schematic diagram illustrating the two RNA synthesis reactions that can occur from the RSV *tr* promoter: *de novo* RNA synthesis, in which RNA is generated by polymerase initiating at the promoter, and back-primed RNA synthesis, in which the *tr* RNA template folds into a secondary structure, with nt 1U and 2G base-pairing with nt 13C and 14A (italicized) and the 3′ end of the RNA is elongated to add in sequence GUC using CAG as a template (underlined). (B) SDS-PAGE analysis of the L-P preparations for the different L mutants. The first lane is a Benchmark ladder. (C) RNA synthesis products generated by the wt and mutant polymerases, analyzed by denaturing polyacrylamide gel electrophoresis. The first two lanes show ladders generated by alkali hydrolysis of an ^32^P end labeled RNA representing the products generated from position 1U and 3C of the promoter, as indicated. (D) Quantification of the data presented in C. The bars show the mean and standard deviation of three independent experiments.

Wt and mutant L-P complexes were expressed by recombinant baculoviruses in insect cells, purified by virtue of a histidine (his) tag on the P protein (Fig 3B) and analyzed for their abilities to engage in *de novo* RNA synthesis from the *tr* promoter and to perform back-priming. This analysis revealed that the P1261A mutant was defective in *de novo* RNA synthesis but generated similar levels of back-primed product as the wt polymerase (Fig 3C and 3D). This indicated that the P1261A mutant had a functional polymerization domain but was defective in synthesizing RNA from the promoter. The W1262A mutant was defective in both *de novo* RNA synthesis and back-priming activities, consistent with the W1262A substitution causing a global polymerase defect (Fig 3C and 3D). These data are consistent with P1261 playing a specific role in RNA synthesis initiation. The W1262 residue could also be involved in initiation, but the fact that it had a global defect in RNA synthesis meant that this could not be easily investigated further, and so subsequent experiments were focused on the P1261A mutant.

### Initiation from position 3C by the P1261A mutant is dependent on initiating NTP concentration

The gel electrophoresis conditions used for the experiments shown in Fig 3 were not appropriate to detect the first products of *de novo* initiation, the pppApC and pppGpA dinucleotides that would be produced from positions 1 and 3, respectively. Therefore, further analysis was performed to determine if the P1261A substitution inhibited initiation (i.e. dinucleotide formation) or the transition to elongation (i.e. formation of products ≥ 3 nt). In addition, we were interested to determine if initiation/ elongation at positions 1 and 3 were affected in the same way. Therefore, we performed experiments to examine synthesis of short products, specifically.

First, we examined RNA synthesis initiation from position 3C of the *tr* promoter. Reactions were performed using a template consisting of nucleotides 1-16 of the *tr* promoter and GTP, ATP and UTP, with [*α*^32^P] ATP included as the radioactive tracer. CTP was omitted to prevent initiation from position 1U (Fig 4A). Under these reaction conditions, it was expected that the wt polymerase would be able to initiate RNA synthesis to form the dinucleotide pppGpA, and then elongate the dinucleotide to the end of the template. As noted in the introduction, a function of a priming residue is to stabilize NTPs within the initiation complex. We reasoned that if Pro1261 fulfills this role, the activity of the P1261A mutant polymerase might depend on the concentration of the initiating NTPs, with higher NTP concentrations increasing the likelihood of them occupying the NTP_1_ and NTP_2_ binding pockets. Therefore, we performed RNA synthesis reactions under conditions in which the initiating GTP and/or ATP were at 10 µM, a concentration which is suboptimal for RNA synthesis initiation, or conditions in which the concentration of ATP and/or GTP was 500 µM, a concentration that we had previously shown to facilitate a high level of initiation from 3C (33). In reactions containing ATP and GTP at 10 µM each, the wt polymerase generated products ranging from 2-14 nt in length, consistent with initiation and elongation from position 3C of the promoter (Fig 4B, lane 3). The identity of the pppGpA dinucleotide was confirmed by omitting either GTP or UTP from the reactions (Fig 4B, lanes 11 and 12). Bands longer than 14 nt were also detected and are most likely due to polymerase stuttering on the U tract, as described previously (14, 34). Examination of products generated by the P1261A mutant showed that under these low NTP concentrations there were no detectable RNA synthesis products, demonstrating that the P1261A substitution inhibits RNA synthesis initiation (Fig 4B, compare lanes 3 and 4, and 4C). A different result was observed in reactions containing 500 µM each of GTP and ATP. In this case, the P1261A mutant was able to generate detectable levels of dinucleotide product, but this product was not elongated (Fig 4B, lane 10). Calculation of the total level of initiation (dinucleotide plus elongation products, normalized for incorporated [*α*^32^P] ATP) showed that with 500 µM GTP and ATP, the P1261A mutant was capable of initiation at 46% of wt levels (Fig 4C). In reactions containing 10 µM GTP and 500 µM ATP, the P1261A mutant had an intermediate phenotype, able to initiate RNA synthesis to a greater degree compared to wt than with 10 µM NTP conditions, but to a lesser degree than with 500 µM conditions (Fig 4B and 4C; note that the band intensities vary for the different NTP conditions, irrespective of levels of RNA synthesis products because the ratios of radioactive to unlabeled ATP in each reaction varied). Together, these results indicate that the P1261A mutant could initiate RNA synthesis at position 3C of the promoter, but that this ability was dependent on the concentration of the initiating NTPs.

**Fig 4.**
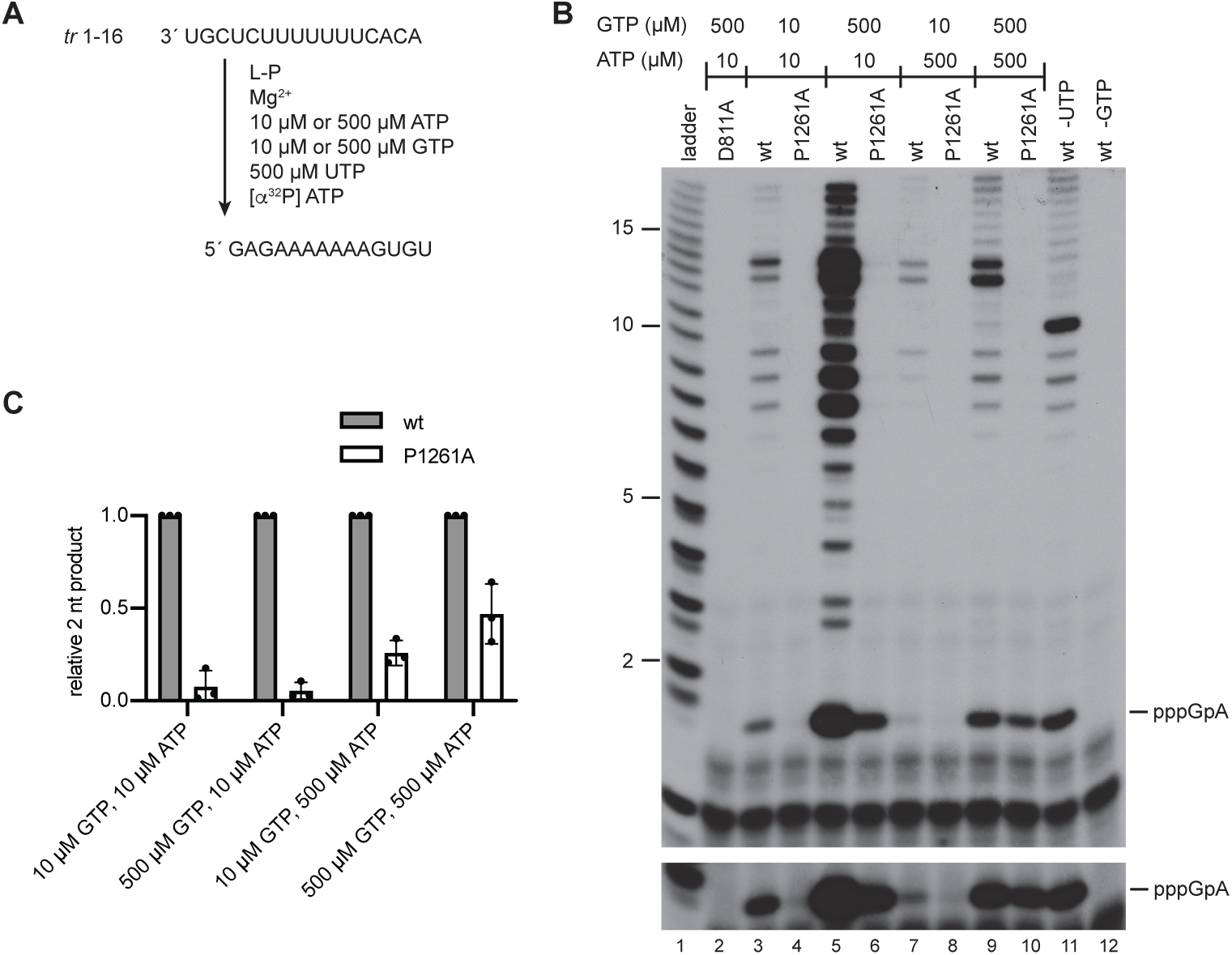
Substitution of P1261A inhibits formation of the initial dinucleotide from the position 3 initiation site, but the defect could be partially rescued with high NTP concentrations. (A) Schematic diagram illustrating the experimental design to detect the initial pppGpA dinucleotide; note that it was not possible to prevent elongation of this dinucleotide. (B) RNA synthesis products generated by the D811A, wt and P1261A polymerase mutants, analyzed by denaturing polyacrylamide gel electrophoresis. Lane 1 shows a ladder generated by alkali hydrolysis of an ^32^P end labeled RNA representing the product generated from position 3C of the promoter. Note that the smaller products of the RSV polymerase migrate slightly further than the corresponding bands in the ladder because they contain a triphosphate rather than monophosphate moiety. The lower panel shows a longer exposure of the image in the upper panel, cropped to highlight the pppGpA products. Note that in this gel, some of the bands appeared as doublets, which presumably represent cleavage intermediates. (C) Quantification of the 2 nt pppGpA band presented in B. The bars show the mean and standard deviation of three independent experiments, normalized to the RNA signal for the wt polymerase in each reaction condition.

### Lack of initiation from position 1U by the RSV L P1261A mutant could not be rescued by increasing initiating NTP concentrations

We then examined pppApC dinucleotide formation from position 1U of the *tr* promoter. Reactions were performed using ATP, CTP, and UTP, with [*α*^32^P] CTP included as the radioactive tracer, and with GTP omitted to prevent initiation from position 3 (Fig 5A). Like the experiment shown in Fig 4, RNA synthesis reactions were performed with suboptimal and optimal concentrations of the initiating NTPs, ATP and CTP. Our previous work has shown that initiation at 1U requires higher concentrations of initiating NTPs than initiation at 3C with the optimal NTP concentrations being 1 mM ATP and 2 mM CTP, and lower concentrations being suboptimal (33). Reactions containing wt polymerase together with 500 µM each of ATP, CTP and UTP yielded products of 2 and 3 nt in length (Fig 5B, lane 7). Given that GTP was omitted from the reactions, a 3 nt product was not expected and so we took steps to determine the identities of the products. Omission of either ATP or UTP from the reaction confirmed that both the 2 and 3 nt products contained ATP and CTP (the radioactive tracer), but not UTP (Fig 5B, lanes 16 and 17). Comparison with a series of di- and tri-nucleotide ladders confirmed that the 2 nt band was the correct size to be pppApC and indicated that the 3 nt band was most likely pppApCpC (Figure 5B, compare lane 7 with lanes 2 and 5). It seems likely that omission of GTP caused the polymerase to engage in non-specific nucleotide addition to generate the pppApCpC product. Although this 3 nt product was an artifact of the reaction conditions, we were still able to assess the ability of the P1261A mutant to initiate RNA synthesis. In contrast to the wt polymerase, the P1261A mutant was defective in dinucleotide (and 3 nt) production under all NTP concentrations that were tested, failing to produce any detectable product above background signal (Fig 5B and 5C). This result is consistent with Pro1261 playing a key role in initiation at position 1U of the template.

**Fig 5.**
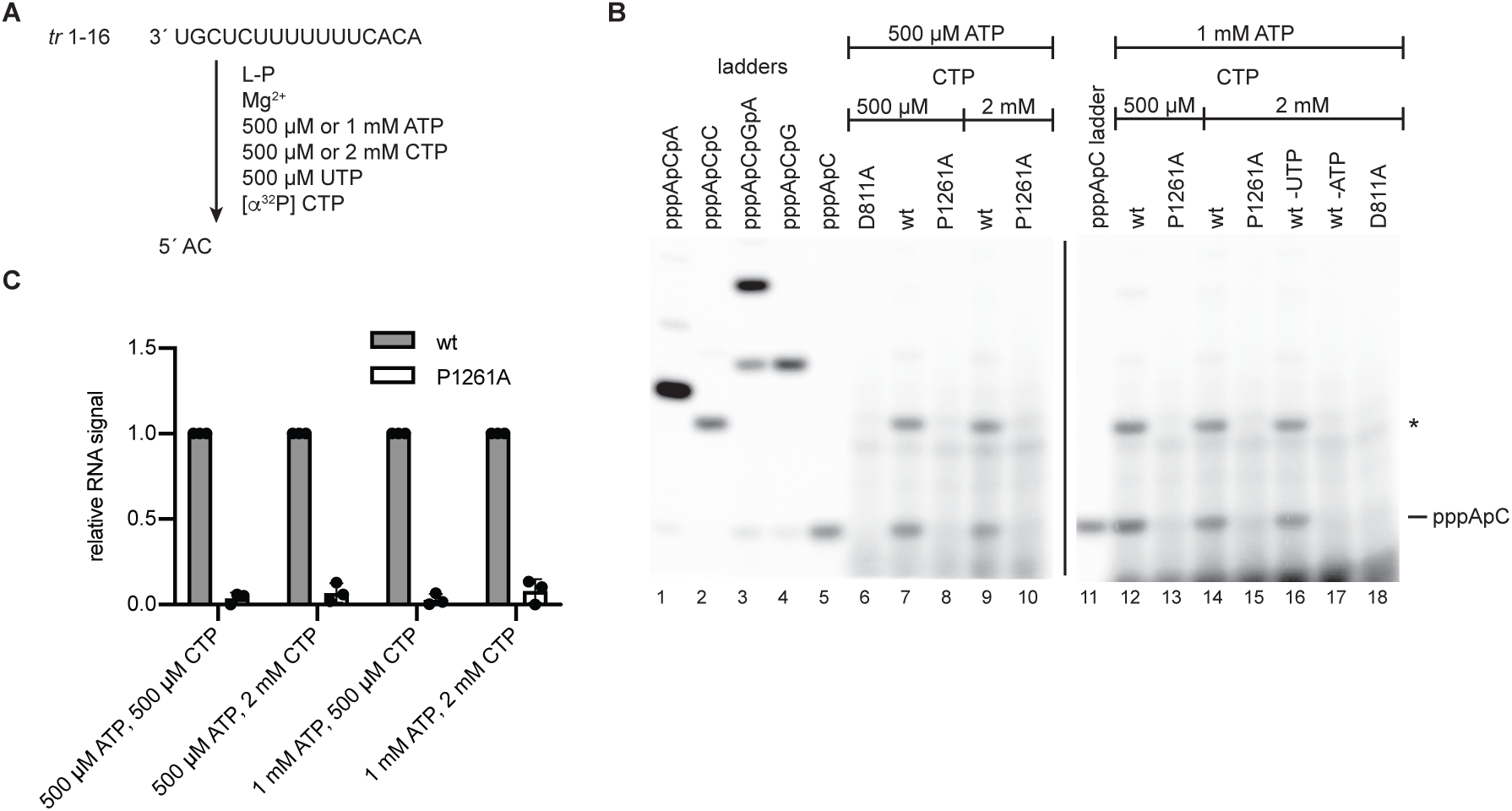
Substitution of P1261A inhibits formation of the initial dinucleotide from the position 1 initiation site, a defect that could not be rescued with optimal NTP concentrations. (A) Schematic diagram illustrating the experimental design to detect the initial pppApC dinucleotide; note that GTP was omitted from the reactions to prevent initiation from position 3. (B) RNA synthesis products generated by the D811A, wt and P1261A polymerase mutants, analyzed by denaturing polyacrylamide gel electrophoresis. Lanes 1-5 show ladders generated by T7 RNA polymerase. The band indicated with an asterisk was an unexpected product, most likely pppApCpC, generated by non-specific terminal transferase activity on the pppApC dinucleotide. (C) Quantification of the data presented in B. The bars show the mean and standard deviation of three independent experiments normalized to the RNA signal for the wt polymerase in each reaction condition. Note that data show the sum of the pppApC and * nt bands, with the band volumes divided by the number of C residues in the product to account for the number of incorporated ^32^P atoms.

### The P1261A defect in initiation could be rescued by replacing Mg^2+^ with Mn^2+^

Polymerases require at least two metal cations for catalysis (35, 36). One metal ion coordinates the 3′ OH of the preceding nucleotide (NTP_1_ during initiation) and the *α*-phosphate of the incoming nucleotide (NTP_2_ during initiation) enabling the nucleophilic attack required for phosphodiester bond formation (37). A second metal ion coordinates the *α*, *β* and *γ* phosphates of the incoming NTP (NTP_2_) helping to neutralize the negative charge and facilitating base-stacking interactions between the nucleobases of the NTPs (37, 38). Although both Mg^2+^ and Mn^2+^ can fulfill these roles, they have different coordination properties: Mn^2+^ binds more tightly to nucleotides than Mg^2+^ and favors initiation by other viral RNA dependent RNA polymerases (37). Consistent with this, Mn^2+^ enhances initiation by the RSV polymerase, but results in a lower level of elongation (34), suggesting that Mn^2+^ stabilizes the initiation complex. Therefore, we investigated if including Mn^2+^ in RNA synthesis reactions could alleviate the initiation defect caused by the P1261A substitution.

Reactions were performed to examine initiation at position 3C of the *tr* promoter under conditions in which either Mg^2+^ or Mn^2+^ was included as the metal cation. Reactions were performed with low concentrations (20 µM) of GTP and ATP, to accentuate defects in initiation. In reactions containing Mg^2+^, the *tr* promoter, GTP and ATP (but no CTP or UTP), the wt polymerase was able to generate a pppGpA dinucleotide and longer products up to 11 nt in length (i.e. up to the first U incorporation site), whereas the P1261A mutant yielded only a very low level of product (Fig 6B, lanes 3 and 4; note that the band intensities for the longer products are lighter than in Fig 4 because the Fig 5 experiment involved radiolabeled GTP, which has fewer incorporation sites than ATP). In contrast, in reactions containing Mn^2+^, both the wt and P1261A polymerases generated equivalent amounts of dinucleotide product (Fig 6A, lanes 6 and 7; Fig 6D). Longer products could not be detected in reactions that contained Mn^2+^, regardless of whether the wt or P1261A polymerase was used. As noted above, the core element of the *tr* promoter is almost identical in sequence to the *le* promoter, but they differ at several positions, including position 4. Given that initiation at position 3C of the *le* promoter requires GTP and CTP (rather than GTP and ATP), we examined if the P1261A substitution affected initiation from this site. For this analysis, we used a *le* promoter containing a G-to-A substitution at position 2, to inhibit initiation at position 1 and allow products initiated at position 3C to be identified using radiolabeled CTP. Like the findings with the *tr* promoter, the P1261A mutant was almost completely defective in initiation from 3C in the presence of Mg^2+^ but initiated RNA synthesis efficiently in the presence of Mn^2+^(Fig 6B and 6E). Together, these data show that the P1261A mutant is conditionally defective in initiation from position 3C, with Mn^2+^ alleviating its initiation defect.

**Fig 6.**
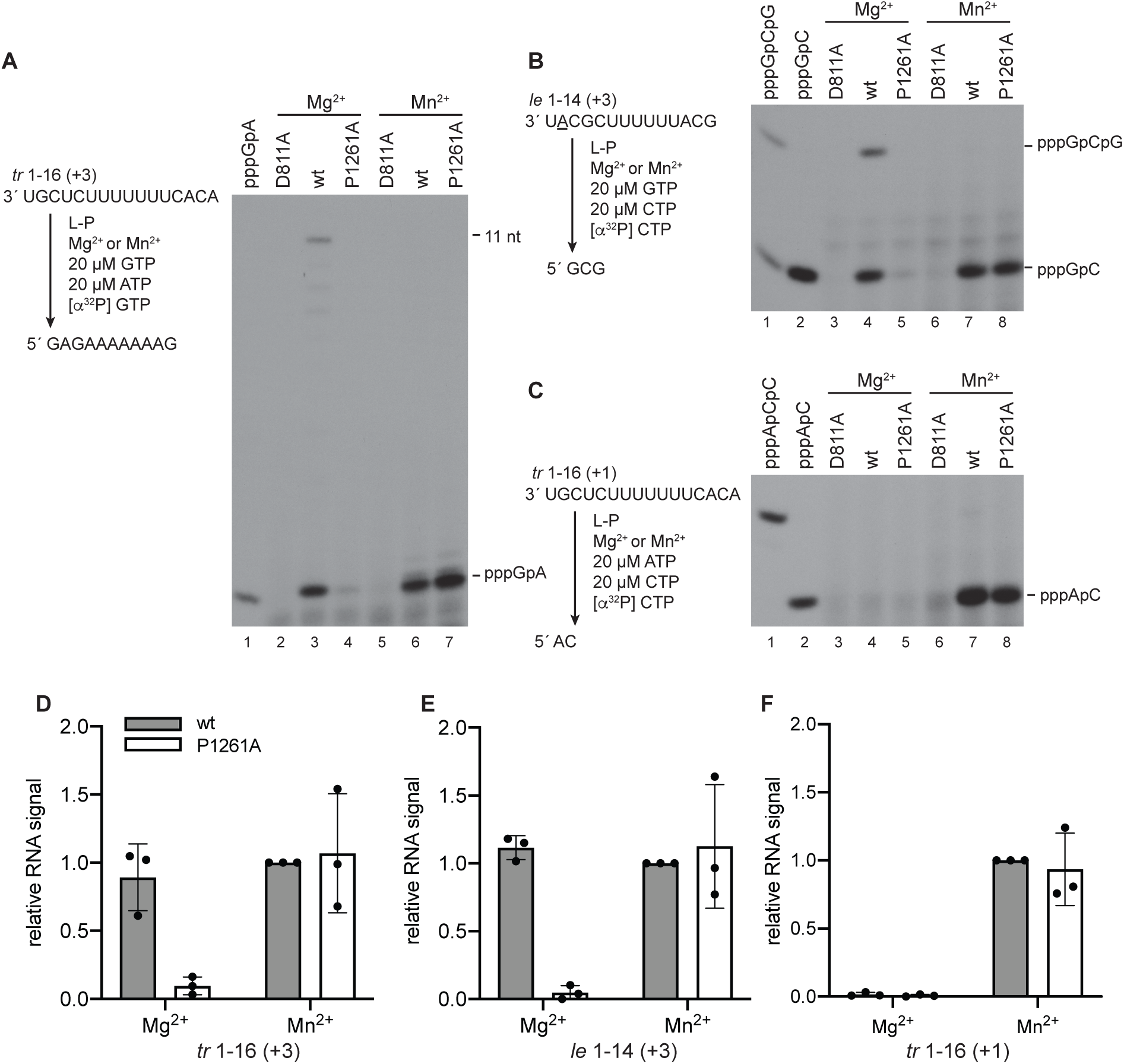
The P1261A initiation defect could be rescued by Mn^2+^. (A-C) RNA synthesis products generated in the presence of Mg^2+^ or Mn^2+^. The left side of each panel shows a schematic diagram illustrating the experimental design and the expected RNA synthesis products in each set of reactions. The site of initiation being analyzed in each reaction is shown in parentheses. In the case of the *le* 1-14 promoter (panel B), a template containing a 2G-to-A substitution (underlined) was used in reactions lacking UTP, to inhibit initiation from position 1. The right side of each panel shows the RNA synthesis products generated by the D811A, wt and P1261A polymerase mutants, analyzed by denaturing polyacrylamide gel electrophoresis. The first lane in panel A and the first two lanes in panels B and C show ladders generated by T7 RNA polymerase. (D, E, F) Quantification of the data presented in A-C. The bars show the mean and standard deviation of three independent experiments, normalized to RNA produced by the wt polymerase in Mn^2+^ conditions. The data show quantification of the sum of the pppGpA and 11 nt bands (D), the sum of the pppGpC and pppGpCpG bands (E) or the 2 nt pppApC band only (F). In D, the band volumes were divided by the number of G residues in the products to account for the number of incorporated ^32^P atoms.

We then examined initiation from position 1U. Because there is only a very low level of initiation from 1U of the *le* promoter, we focused our analysis on the *tr* promoter. Reactions were performed with 20 µM ATP and CTP, as described above. Under these limiting NTP concentrations, reactions that contained Mg^2+^ did not yield detectable pppApC dinucleotide from position 1 of the *tr* promoter, with either wt L-P or the P1261A mutant (Fig 6C, lanes 4 and 5). However, in reactions containing Mn^2+^, a significant amount of product was generated by both polymerases (Fig 6C, lanes 7 and 8; Fig 6F). These results show that Mn^2+^ significantly enhanced initiation by the wt polymerase, even under these very low NTP concentrations, and completely restored the P1261A defect.

### The residues of MARV L protein corresponding to RSV L Pro1261 and Trp1262 are not required for RNA synthesis initiation

As shown in Fig 1B, a proline residue at a position corresponding to the RSV L Pro1261 is conserved in many, but not all nsNSVs. This raised the question as to whether the function of this proline residue is conserved in the virus polymerases in which it is present. As shown in the alignment presented in Fig 1B, the filovirus MARV L protein does not have an aromatic residue corresponding to the rabies virus L Trp1180, but it does have a proline residue (Pro1217) that aligns with RSV L Pro1261 and an aromatic tyrosine residue (Y1218) that aligns with RSV L Trp1262. Since we have recently established an *in vitro* RNA synthesis initiation assay for the MARV polymerase (39), we used this approach to test the effect of substituting MARV L Pro1217 and Tyr1218 with alanine. The mutant L-VP35 complexes were isolated and tested in an on-bead RNA synthesis assay using an RNA template consisting of nucleotides 1-30 of the MARV *le* promoter sequence (Fig 7A and 7B). As a negative control, an L mutant with a substitution in the aspartic acid residue of the GDNQ RNA dependent polymerase motif was generated (L D774A). Surprisingly, we found that both MARV L P1217A and Y1218A mutants efficiently synthesized RNA, compared to the wt L protein (Fig 7C). Thus, although Pro1261 plays a key role in RSV RNA synthesis initiation and Trp1262 is required for RSV RNA synthesis, the corresponding residues of the MARV L protein do not play similar roles.

**Fig 7.**
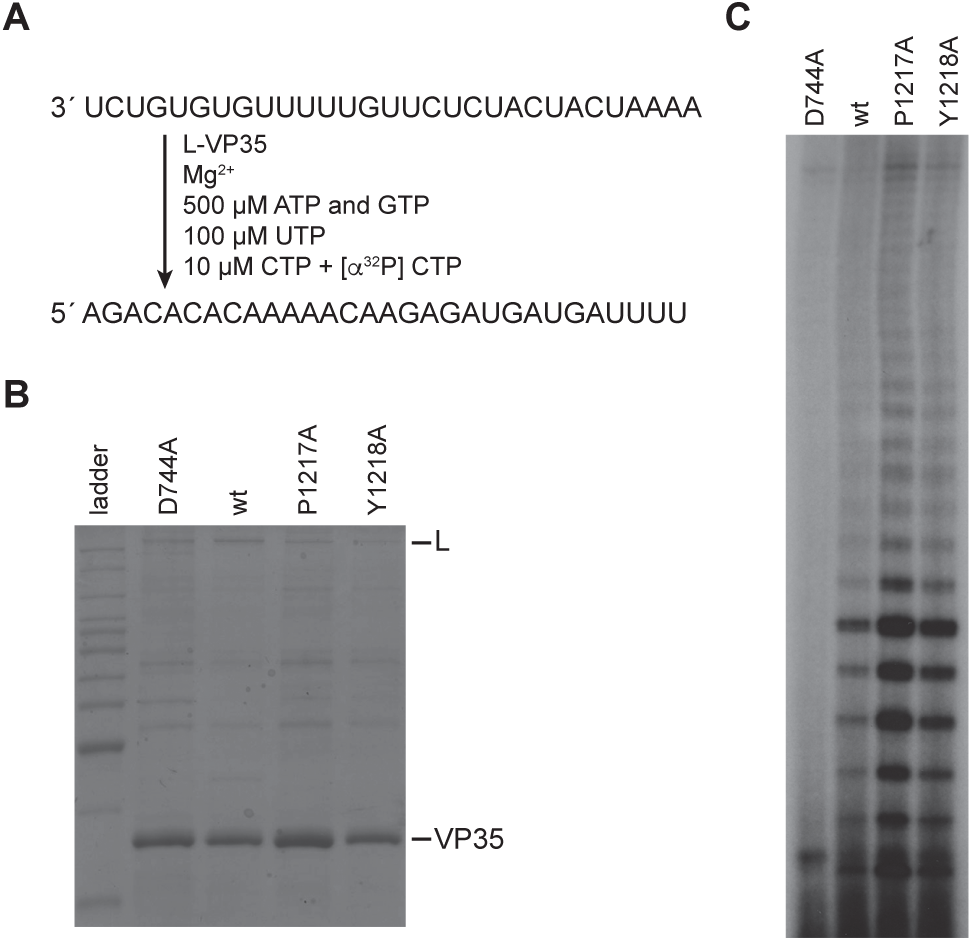
MARV L Pro1217 and Tyr1218 are not required for RNA synthesis initiation. (A) Schematic diagram showing the sequence of the MARV *le* 1-30 promoter template, the reaction conditions used to assess RNA synthesis, and the expected product RNA. (B) SDS-PAGE analysis of the L-VP35 preparations for the different L mutants. The first lane is a Benchmark ladder. (C) RNA synthesis products generated by the wt and mutant polymerases, analyzed by denaturing polyacrylamide gel electrophoresis.

## Discussion

The transcription-replication complexes of the nsNSVs share many features. However, at a detailed level, there are differences between them. For example, whereas rhabdoviruses, paramyxoviruses and filoviruses have a single initiation site in their promoters, the pneumoviruses, RSV and HMPV, have two initiation sites (9, 14, 30, 39, 40) and whereas as most nsNSVs investigated so far initiate RNA replication at position 1 of the promoter, ebolaviruses initiate at position 2 (41, 42). It is of interest to understand to what extent these different mechanisms reflect differences in the polymerase protein. In a previous study, we showed that although the PIV-3 polymerase initiates only at position 1 of its own promoter, it initiates at positions 1 and 3 of the RSV promoter (like the RSV polymerase). Likewise, we showed that although MARV polymerase only initiates at position 1 of its own promoter, it too can initiate efficiently at an internal site at position 2 of the Ebola virus promoter (39). These findings indicate that while nsNSVs have differences in their promoters and sites of initiation, their polymerases have similar properties. Given that available structures of pneumovirus and paramyxovirus polymerases show that they contain a loop structure in a similar position as the rhabdovirus priming loop, albeit in a different conformation, it was reasonable to consider that ring-based residues in this region might play a role in initiation of RNA synthesis by other nsNSVs. The goal of this study was to investigate the priming loop feature of the RSV polymerase and specifically to ascertain if conserved proline and tryptophan residues were involved in RNA synthesis initiation. We also sought to determine if these conserved residues play an equivalent role in the MARV polymerase.

In a previous study, we showed that substitution of Pro1261 or Trp1262 completely inhibited initiation and/or early elongation from positions 1 and 3 of the *tr* promoter (30). Here we showed that the P1261A and W1262A mutants were defective in initiation from positions 1 and 3 of the *le* promoter, although in this case the 3C defect was not as severe (Fig 2). This suggested that Pro1261 and/ or Trp1262 might function to stabilize the initiation complex, in a manner similar to a priming residue. While study of the W1262A mutant was hampered by the fact that it appeared to have a general defect in RNA synthesis, analysis of the P1261A substitution showed that it inhibited initiation without inhibiting back-priming (Fig 3). This indicates that the P1261A substitution caused specific initiation defects rather than general dysfunction due to aberrant folding of the polymerase. We examined the function of the P1261A mutant under conditions that could be expected to increase initiation complex stability, including increased concentration of the initiating NTPs and including Mn^2+^ rather than Mg^2+^ as the metal cation. In both cases, we found that the P1261A defect was ameliorated to some extent by the change in conditions, consistent with it having a role in RNA synthesis initiation. The impact of each of these factors and promoter sequence is discussed below.

Increasing the concentration of GTP and ATP, the initiating NTPs for 3C initiation, from 10 to 500 mM increased the relative 3C initiation efficiency of the P1261A mutant from ∼10 to 50% (Fig 4), consistent with the possibility that Pro1261 stabilizes the initiating NTPs. Initiation at 1U was not rescued with increased NTP concentrations (Fig 5), however, this might reflect different Km values for the initiating NTPs for the two different initiation events as higher concentrations of initiating NTPs are required for initiation at 1U than 3C (33). Therefore, it is possible that even the relatively high concentrations of 1 mM ATP and 2 mM CTP were insufficient to rescue the P1261A defect in 1U initiation. As noted in the results section, metal cations interact with NTPs to facilitate polymerization and Mn^2+^ has a higher affinity for NTPs than Mg^2+^. In the experiments presented here, Mn^2+^ had no effect on the ability of the wt polymerase to initiate RNA synthesis at position 3C, but completely rescued the 3C initiation defect of the P1261A mutant, even with low NTP concentrations. Again, this is consistent with Pro1261 playing a role in stabilizing NTPs within the initiation complex. Mn^2+^ had an intriguing effect on initiation from 1U, enabling even the wt polymerase to initiate with much greater efficiency at position 1U (Fig 6). This suggests that Mn^2+^ helped stabilize the wt polymerase 1U initiation complex even though it was not necessary to stabilize the wt polymerase 3C initiation complex. This again is consistent with the concept that there is a higher Km value for the 1U versus the 3C initiating NTPs. Mn^2+^ also completely alleviated the 1U initiation defect caused by the P1261A substitution (Fig 6), consistent with this proline residue affecting the stability of 1U initiation complex, in addition to the 3C initiation complex.

We also found that the effect of the P1261A substitution was different for the *le* versus *tr* promoter (Fig 2). In experiments with the *tr* promoter the P1261A substitution completely inhibited detectable RNA synthesis from both 1U and 3C (30), whereas on the *le* promoter, initiation from 1U was completely inhibited, but initiation from 3C occurred at ∼25% of wt levels (Fig 2). This finding indicates that Pro1261 is required to a greater extent for 3C initiation on the *tr* promoter than on the *le* promoter. The difference at position 4 of the *le* and *tr* promoters means that their respective position 3 initiation complexes involve different initiating NTPs (12, 40). Our previous studies have shown that initiation from position 3C is considerably more efficient on the *le* compared to the *tr* promoter, suggesting that the *le* position 3C initiation complex is more energetically stable (33). Given that the first 9 residues are sufficient for efficient promoter activity (43), and that the *le* and *tr* promoters are identical for 10 of the first 11 nt, it is likely that the different ratios of 1U: 3C initiation are due to a difference at position 4 of the promoters, which results in different initiating NTPs being used for 3C initiation. In the *le* 3C initiation complex, NTP_2_ is a C residue, capable of having three hydrogen bonds in Watson-Crick base-pairing with the template, whereas in the *tr*, NTP_2_ is an A residue, capable of only two hydrogen bonds. This differential in hydrogen bonding potential could allow the *le* 3C initiation complex to have greater stability than the *tr* 3C initiation complex, allowing the RNA synthesis defect caused by the P1261A substitution to be overcome to some extent. Thus, this result is also consistent with Pro1261 being involved in stabilizing the initiating NTPs.

Taken together, these data suggest that Pro1261 is involved in stabilizing the initiating NTPs for both 1U and 3C initiation. In the RSV polymerase structure, Pro1261 and Trp1262 are in the middle of a loop region that aligns with the priming loop sequence of VSV and rabies virus, but which is folded against the capping domain, out of the active site of the polymerization domain (10, 26) (Fig 1A). Proline has been shown to stack with tryptophan in a peptide sequence (44) and a structural simulation using ISOLDE suggests that it is physically possible for either Trp1262 or Tyr 1276 to act as a priming residue. If Tyr1276 acts as the priming residue, then the hinge point for the RSV L priming loop will resemble the hinge of the rhabdovirus L priming loops (Fig 8A-B). If Trp1262 acts as the priming residue, then the priming loop hinge on RSV L will differ from rhabdovirus L priming loops (Fig 8A, Fig 8C). Based on the data presented here, we propose that Pro1261 and Trp1262 cooperate to form base-stacking interactions with the initiating NTPs to facilitate initiation. We also propose that Pro1261 has greater significance for initiation at 1U than 3C, due to a high Km required for the 1U initiating NTPs. An alternative explanation is that Pro1261 plays a structural role and that inclusion of Mn^2+^ and promoter sequence differences cause polymerase structural changes that allow the P1261A mutant to overcome its defect. A structure of the polymerase captured as an initiation complex is likely to be required to distinguish between these possibilities.

**Fig 8.**
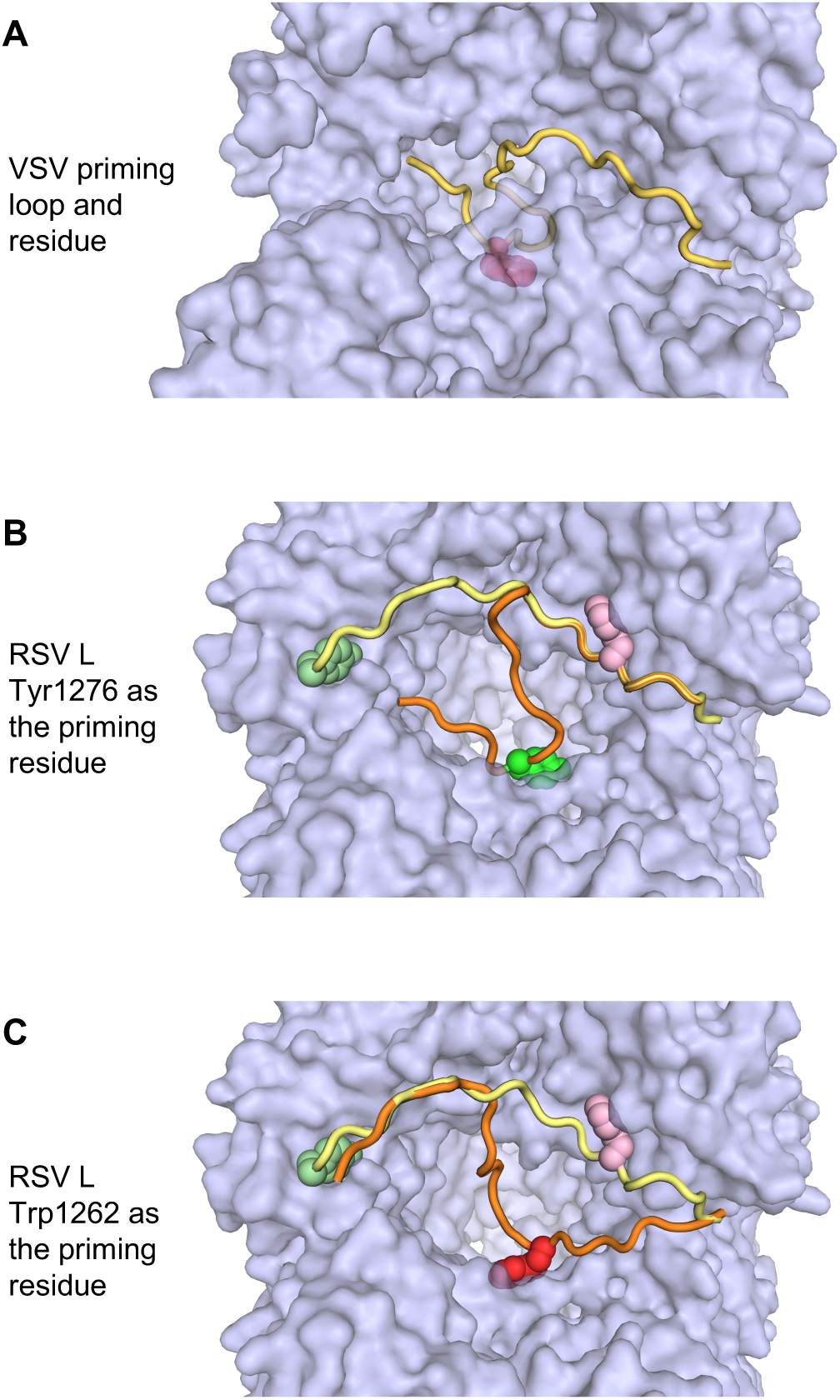
Structure simulation indicates that RSV L Trp1262 could function as a priming residue. (A) Space filling model of the VSV polymerase showing the RNA synthesis active site and priming residue. The priming loop is shown in gold with the priming tryptophan residue in magenta. (B and C) Models of the RSV polymerase showing the priming loop in different conformations. The priming loop is shown in both its retracted state (yellow), as in the solved structures, or protruding into the active site cavity (orange), according to the simulation. Tyr1276 is shown in different shades of green, and Trp1262 in pink or red, depending on their positioning. The ISOLDE simulation shows that either Tyr1276 or Trp1262 is capable of protruding into the active site (panels B and C, respectively).

In addition to its role in initiation Pro1261 also seems to affect elongation, at least in *in vitro* reactions, inhibiting efficient elongation beyond a 2 nt product from the 3C initiation site of the *tr* promoter, for example (Fig 4). However, this elongation inhibition was not absolute as the P1261A mutant could generate elongated RNA from the 3C initiation site of the *le* promoter in the minigenome system, albeit at a low level (Fig 2). It is possible that elongation is affected by promoter sequence and/or the local environment in transfected cells, with factors such as the N-RNA template perhaps playing a role to overcome an elongation defect (45).

Regardless of the mechanism by which the P1261A substitution affects RSV polymerase function, substitution at this site is clearly highly deleterious to RNA synthesis initiation. Sequence alignments indicated that this proline residue is conserved across many (although not all) nsNSVs (Fig 1B), suggesting that it might play a similar function in other viruses. However, alanine substitution of the corresponding proline residue of the MARV L protein (Pro1217) was well tolerated for the process of RNA synthesis initiation, as was substitution of the adjacent aromatic Tyr1218 residue (Fig 7). This combined with the finding that the rhabdovirus tryptophan priming residue is not conserved across nsNSVs shows that there are differences between nsNSV polymerases, even in conserved motifs. Similar observations have been made regarding the conserved GxxT motif and its role in capping (5). Thus, even though there are elements of sequence, structure and functional conservation in the polymerases of nsNSV families, evidence is accumulating to indicate that they have evolved features that distinguish them from each other.

## Materials and Methods

### L plasmid mutagenesis

Mutations were introduced into a codon optimized version of the L ORF of RSV strain A2. Mutations in the GDNQ motif of the polymerization domain (D811A), the priming loop/ capping domain (P1261A, W1262A, P1274A, and Y1276A) have been described previously (5, 30). Briefly, mutations were generated in a fragment of the L ORF subcloned into a pGEM T easy vector (Promega). The mutated ORF fragment was sequenced and then substituted in the L ORF of pTM1, a T7 expression vector in which the ORF lies immediately following an internal ribosome entry site, or assembled into an L ORF contained in a pFastBac Dual vector (Invitrogen) using the NEBuilder HiFi DNA Assembly Cloning kit. The pFastBac Dual vector also contained the RSV A2 P ORF tagged with the tobacco etch virus (TEV) protease cleavage site and a hexahistidine sequence, as described previously (14). Bacmids and recombinant baculoviruses were generated from the pFastBac Dual vectors using the Bac-to-Bac system (Invitrogen), according to the manufacturer’s instructions. To examine expression of the mutant L proteins in mammalian cells, the pTM1 plasmid containing wt L was modified to insert two tandem FLAG tags at the L N terminus by insertion of an oligo duplex. L fragments containing the priming loop mutations were then inserted into this plasmid. All DNA clones were sequenced to confirm the presence of mutations, and to ensure that no other changes were introduced during PCR-mediated mutagenesis.

### Reconstitution of minigenome replication and transcription in cells

Polymerase activities were assessed in the minigenome system using pTM1 vectors containing untagged versions of the L proteins. The minigenome template that was used has been described previously (9). It was a dicistronic minigenome containing the 44 nt *le* promoter, *NS1 gs* sequence and *NS1* non-translated region at the 3′ end and *L gene end* signal and trailer region at the 5′ end. The minigenome was limited to the antigenome step of replication due to a substitution at position 2 relative to the 5′ end of the trailer region, which inactivated the *tr* promoter at the 3′ end of the antigenome. Minigenome RNA synthesis was reconstituted in BSR-T7/5 cells, as described previously (46), except that the cells were incubated at either 30°C or 37°C, as indicated. At 48 h post-transfection, cells were harvested to isolate total intracellular RNA. RNA samples were prepared using Trizol (Invitrogen) according to the manufacturer’s instructions, except that following the isopropanol precipitation step, the RNA was subjected to additional purification by acid phenol-chloroform extraction and ethanol precipitation.

### Primer extension analysis of minigenome derived RNAs

Primer extension reactions were performed on total intracellular RNA isolated from minigenome transfected cells. Analysis of RNA initiated at the +1 and +3 sites of the *le* promoter was performed with reactions containing 0.2 µM radiolabeled primer that hybridized to nucleotides 15-39 relative to the 5′ terminus of the antigenome RNA (5′ TTTGGTTTATGCAAGTTTGTTGTAC), 500 µM dNTPs, and Sensiscript reverse transcriptase (Qiagen), following the manufacturer’s instructions. These primer extension products were compared to ^32^P-end labeled DNA oligonucleotides of sequence and length equivalent to cDNAs corresponding to RNAs initiated at positions 1U and 3C. Analysis of RNA initiated at the *gs* signal was performed with reactions containing 10 nM radiolabeled primer that hybridized to nucleotides 12-31 relative to the 5′ end of the templated mRNA (5′ AAGTGGTACTTATCAAATTC), 2 µM dNTPs and MMLV reverse transcriptase (Promega). Primer extension reactions to analyze initiation from the +1 and +3 sites of the *le* promoter were subjected to electrophoresis in 6% polyacrylamide gels containing 7 M urea in Tris-borate-EDTA buffer. Primer extension reactions to analyze initiation from the *gs* signal were subjected to electrophoresis in 20% polyacrylamide gels containing 7 M urea in Tris-borate-EDTA buffer. Gels were dried onto 3MM paper using a vacuum drier at 80°C. Primer extension products were detected by autoradiography and quantified by phosphorimager analysis.

### Purification of the RSV L-P and MARV L-VP35 complexes

Sf21 insect cells cultured in suspension in SF-900 II serum free medium (Invitrogen) were infected with recombinant baculovirus expressing either wt or mutant RSV or MARV L protein together with either RSV P or MARV VP35 containing a C-terminal histidine tag. RSV L-P protein complexes were isolated from cell lysates by affinity chromatography on Ni-NTA agarose resin (ThermoFisher) and MARV L-VP35 complexes were isolated no Ni-NTA magnetic beads (ThermoFisher). All lysis, wash, and elution buffers contained 50 mM NaH_2_PO_4_ (Sigma), 150 mM NaCl (Sigma), 0.5% NP40 (IGEPAL CA-630, Sigma), pH 8.0 and the indicated concentration of imidazole. Cells were lysed in 20 mM imidazole containing buffer. For the RSV L-P complexes, the resin was washed three times with 60 mM imidazole containing buffer, two times with 100 mM imidazole containing buffer, and the L-P complex was eluted with 250 mM imidazole containing buffer. The L-P preparation was then dialyzed against 150 mM NaCl, 20 mM Tris-HCl pH 7.4 (Sigma), 10% glycerol dialysis buffer. For the MARV L-VP35 complexes, the magnetic beads were washed two times with 100 mM imidazole containing buffer and two times with 200 mM imidazole containing buffer. The MARV L-VP35 protein complex was stored retained on the magnetic beads in the same storage buffer as the RSV L-P complex. Isolated L-P and L-VP35 complexes were analyzed by SDS-PAGE and PageBlue staining (Fermentas) and the L protein concentration was estimated by comparing its band intensity to bovine serum albumin reference standards.

### In vitro RNA synthesis reactions to detect RSV polymerase products

Reactions with *tr* 1-25 templates and [*α*-^32^P]-GTP tracer were performed in 25 μL reactions with the following conditions: 2 μM RNA oligonucleotide (Dharmacon), 50 mM Tris, pH 7.4, 8 mM MgCl_2_, 5 mM DTT, 10% glycerol, 500 μM (each) ATP, CTP, GTP, and UTP with 5 μCi [*α*-^32^P]- GTP (3000 Ci/mmol). Reaction mixtures were preincubated at 30°C for 5 minutes prior to addition of L-P complexes to a concentration of 8 nM L protein. Reaction mixes were incubated at 30°C for 1 h and then heat inactivated at 90°C for 3 minutes then allowed to cool on ice for 2 minutes. RNA was extracted with phenol-chloroform and ethanol precipitated. Pellets were resuspended in RNase free water, and an equal volume of stop buffer (deionized formamide containing 20 mM EDTA, bromophenol blue, xylene cyanol) was added. Products were migrated alongside molecular weight ladders representing products initiated from 1U or 3C sites, prepared as described previously (14). RNA samples were subjected to electrophoresis on a 20% polyacrylamide gel containing 7 M urea in Tris-borate-EDTA buffer. Gels were dried onto 3MM paper using a vacuum drier at 80°C and analyzed by autoradiography and phosphorimage analysis.

### In vitro RNA synthesis reactions to detect dinucleotide formation under varying NTP conditions

Reactions to detect pppApC formation from position 1 of the promoter were performed in a volume of 25 μL with the following conditions: 2 μM *tr*1-16 RNA oligonucleotide (Dharmacon), 50 mM Tris, pH 7.4, 8 mM MgCl_2_, 5 mM DTT, 10% glycerol, 500 μM or 1 mM ATP, 500 μM or 2 mM CTP and 100 μM UTP with 10 μCi [*α*-^32^P]-CTP (3000 Ci/mmol). Reactions to detect pppGpA and elongation products generated from position 3 of the promoter were performed in a volume of 25 μL with the following conditions: 2 μM RNA oligonucleotide (Dharmacon), 50 mM Tris, pH 7.4, 8 mM MgCl_2_, 5 mM DTT, 10% glycerol, 10 or 500 μM ATP, 10 or 500 μM GTP, and 500 μM UTP with 5 μCi [*α*- ^32^P]-ATP (3000 Ci/mmol). In all cases, reaction mixtures were preincubated at 30°C for 5 minutes prior to addition of L-P complexes to a concentration of 8 pM L protein. Reaction mixes were incubated at 30°C for 1 h and then heat inactivated at 90°C for 3 minutes then allowed to cool on ice for 2 minutes. Following inactivation of the polymerase, all reactions were adjusted to contain 500 μM GTP (this avoided the loss of small RNA products that occurred during subsequent purification) and RNA was extracted with phenol-chloroform and ethanol precipitated. Pellets were resuspended in RNase free water, and an equal volume of stop buffer (deionized formamide containing 20 mM EDTA, bromophenol blue, xylene cyanol) was added. The samples were subjected to denaturing electrophoresis alongside T7 triphosphorylated RNA markers by electrophoresis on 25% polyacrylamide gels containing 7 M urea in Tris-taurine-EDTA buffer. Gels were analyzed directly (without drying) by autoradiography and phosphorimage analysis.

### In vitro RNA synthesis reactions to detect dinucleotide formation with Mg^2+^ versus Mn^2+^ conditions

Reactions were performed in a volume of 25 μL with the following conditions: 2 μM RNA oligonucleotide (Dharmacon), 50 mM Tris, pH 7.4, 8 mM MnCl_2_ or MgCl_2_, 5 mM DTT, 10% glycerol. The pppGpC reactions contained *le* 1-14 2G/A RNA template, 20 µM (each) GTP and CTP, and 5 µCi [*α*^32^P]-CTP (3000 Ci/mmol) as the radioactive tracer. The pppApC reactions contained *tr* 1-16 RNA template, 20 µM (each) ATP and CTP, and 5 µCi [*α*^32^P]-CTP (3000 Ci/mmol) as the radioactive tracer. The pppGpA reactions contained *tr* 1-16 RNA template, 20 µM (each) GTP and ATP, and 5 µCi [*α*^32^P]-GTP (3000 Ci/mmol) as the radioactive tracer. Reaction mixes were preincubated prior to addition of L-P protein to a concentration of 8 pM L protein. Reaction mixes were incubated at 30°C for 1 h, heat inactivated at 95°C for 5 minutes, and allowed to cool on ice for 2 minutes. Reactions were adjusted to contain 500 μM GTP after inactivation of the polymerase and RNA was purified and analyzed as described above.

### Synthesis of 5′ triphosphorylated RNA markers

Markers consisting of 5′ triphosphorylated RNA used to characterize small RNA products were generated using a method adapted from (47). DNA oligonucleotides (shown in Table 1) were annealed to create an oligo duplex of the Class II T7 promoter, and a template for synthesis of a small 5′ triphosphorylated RNA by T7 *in vitro* transcription. Annealed oligonucleotides were prepared by mixing 4 μM short oligonucleotide and 4 μM long overhang (template) oligonucleotide in Tris-EDTA pH 8.0, incubated at 90°C for 3 minutes, then allowed to cool to room temperature. 40 μL *in vitro* transcription reactions were prepared with the following conditions: 40 mM Tris-HCl (pH 8.0), 20 mM MgCl_2_, 2 mM spermidine,10 mM DTT, 0.1 mg/mL bovine serum albumin, 2 mM ATP, 1 mM CTP, 1 mM GTP with 1 μCi [α-^32^P] ATP, [α-^32^P] CTP, or [α-^32^P] GTP depending on the marker being synthesized, 0.1 μM DNA annealed oligos and 1 unit/μL T7 RNA polymerase (New England Biolabs). Reactions were incubated at 37°C for 5 hours. Samples were treated with 1 μL DNase I (New England Biolabs) for 10 minutes at 37°C. 1 μL 0.5 M EDTA was added to the reactions. Products were subjected to phenol chloroform extraction and ethanol precipitation prior to use as markers.

**Table 1.**
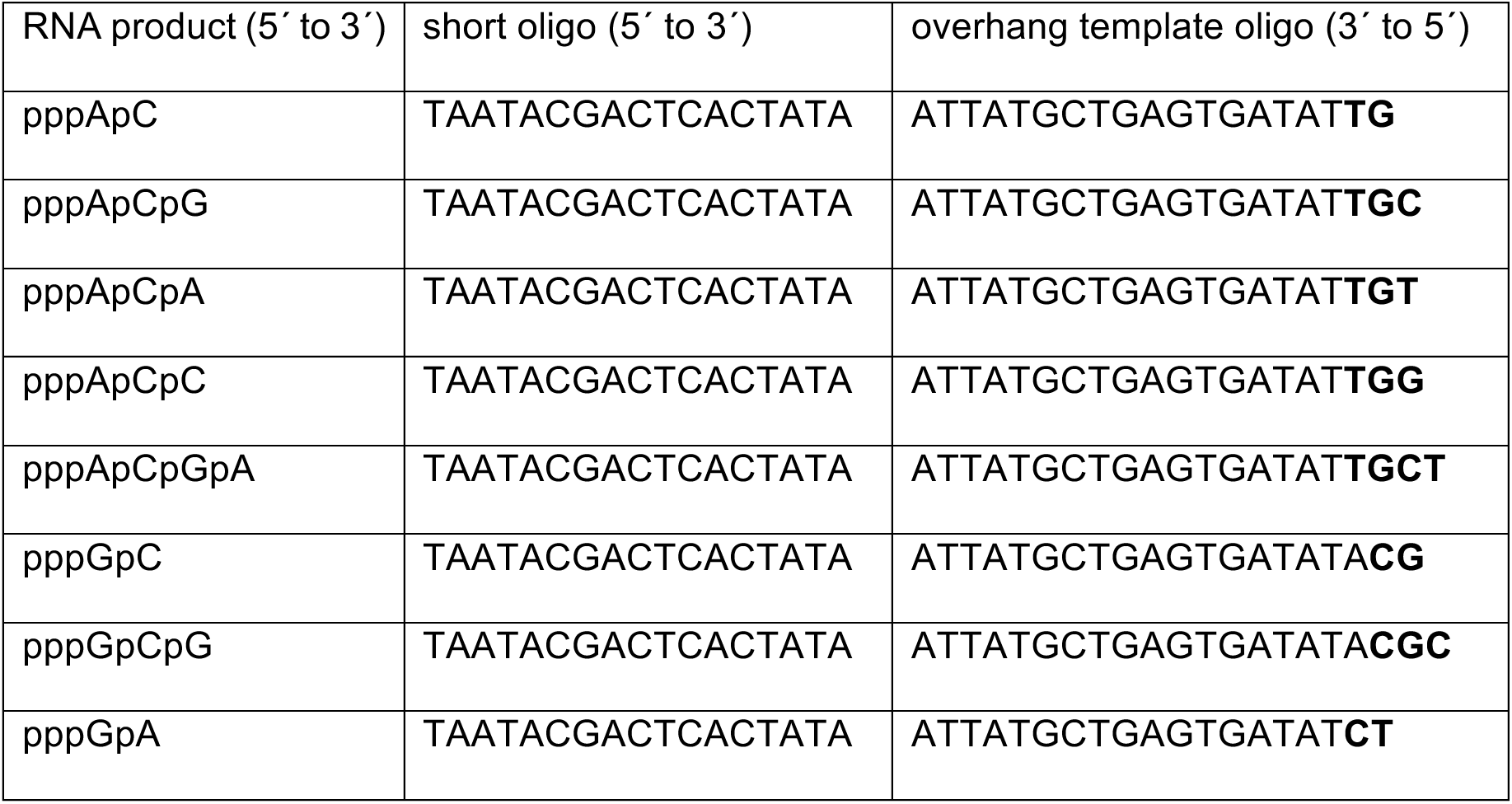
DNA oligonucleotides used for T7 in vitro transcription synthesis of 5′ triphosphorylated RNA markers. Templating sequence is in bold type.

### In vitro RNA synthesis reactions to detect MARV polymerase products

Reactions were performed in a volume of 25 μL with the following conditions: 2 μM MARV *le* 1-30 RNA oligonucleotide (Dharmacon), 50 mM Tris, pH 7.4, 8 mM MgCl_2_, 5 mM DTT, 10% glycerol, 500 μM (each) ATP and GTP, 10 µM CTP, and 100 μM UTP, with 10 μCi [a-^32^P]-CTP (3000 Ci/mmol) as the radioactive tracer. Reaction mixes were preincubated prior to addition of L-VP35 complexes to a concentration of 8 nM L protein. Reaction mixes were incubated at 30°C for 1 h, heat inactivated at 95°C for 5 minutes, and allowed to cool on ice for 2 minutes. RNA was purified and analyzed on 25% Tris-taurine gels, as described above.

### Confirmation of mutant L protein expression in mammalian cells

To confirm that each of the mutant L proteins could be expressed in mammalian cells, BSR-T7/5 cells were transfected with pTM1 plasmids containing FLAG-tagged versions of wt or mutant L plasmids. Cells in a 6-well plate were transfected with 2 μg of pTM1 P and 1 μg wt or mutant L using Lipofectamine 2000 (Invitrogen), according to the manufacturer’s instructions. The cells were incubated at 30°C or 37°C (as indicated) and at 48 h post-transfection, they were collected and incubated with lysis buffer (150 mM NaCl, 0.5% NP40, 50 mM NaH_2_PO_4_, 20 mM imidazole) for 15 minutes on ice, then treated with Turbo DNase (Invitrogen) for 5 minutes and then migrated on a 10% SDS polyacrylamide gel and analyzed by Western blot analysis using a monoclonal antibody specific for the FLAG tag (Sigma). Proteins were detected using a goat anti-mouse IgG coupled to IRDye 800CW and LI-COR analysis.

### Quantification of RNA products and image presentation

Primer extension products and RNA products generated in biochemical assays were quantified by phosphorimage analysis. In all cases, the value obtained from a corresponding region of a negative control lane was subtracted from each of the values of the test lanes. In the case of the primer extension analysis, the data were normalized to the level of input minigenome, as determined by Northern blot analysis, to control for variation in transfection efficiency and RNA purification. In some of the images presented, the contrast was adjusted using the brightness/ contrast function in Adobe Photoshop. Any adjustments made were applied to the entire image.

### Molecular Modeling of Priming Loops

Models for the RSV L priming loop were built manually in Coot (48), then imported into ChimeraX (49) and minimized using ISOLDE (50) to eliminate steric clashes and optimize model geometry.

## Acknowledgements

We thank Dr. Barbara Ludeke for critical reading of the manuscript.

---

**S1 Fig.**
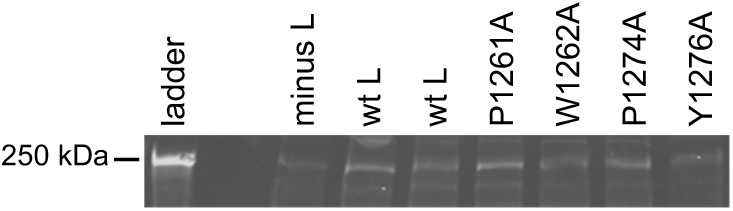
Analysis of L mutant protein expression in BSR-T7 cells at 37°C. To quantify mutant L protein levels in transfected cells, the proteins were modified by adding two tandem FLAG tags to their N-termini. The FLAG-tagged L mutants were co-expressed with P protein in BSRT7/5 cells. At 48 h post transfection, cell lysates were analyzed by Western blotting using a FLAG-specific antibody. The image shown is representative of three independent experiments. Note that a much larger amount of L plasmid was required for protein detection by Western blotting than is optimal for minigenome assays and so we were unable to analyze protein levels in the same cells that were used for the minigenome RNA analysis shown in Fig 2, which employed lower levels of untagged L protein. A background band (visible in the minus L lane) complicated the analysis somewhat, but nonetheless, the results showed that all mutant proteins were expressed. The levels of the W1262A L protein were variable, and it was sometimes expressed at a lower level than that of the other mutants, suggesting that it might have been unstable at this temperature. However, we found that it was stable at 30°C and the levels of RNA synthesis for each of the L mutants at 30°C was similar as at 37°C (see S2 Fig).

**S2 Fig.**
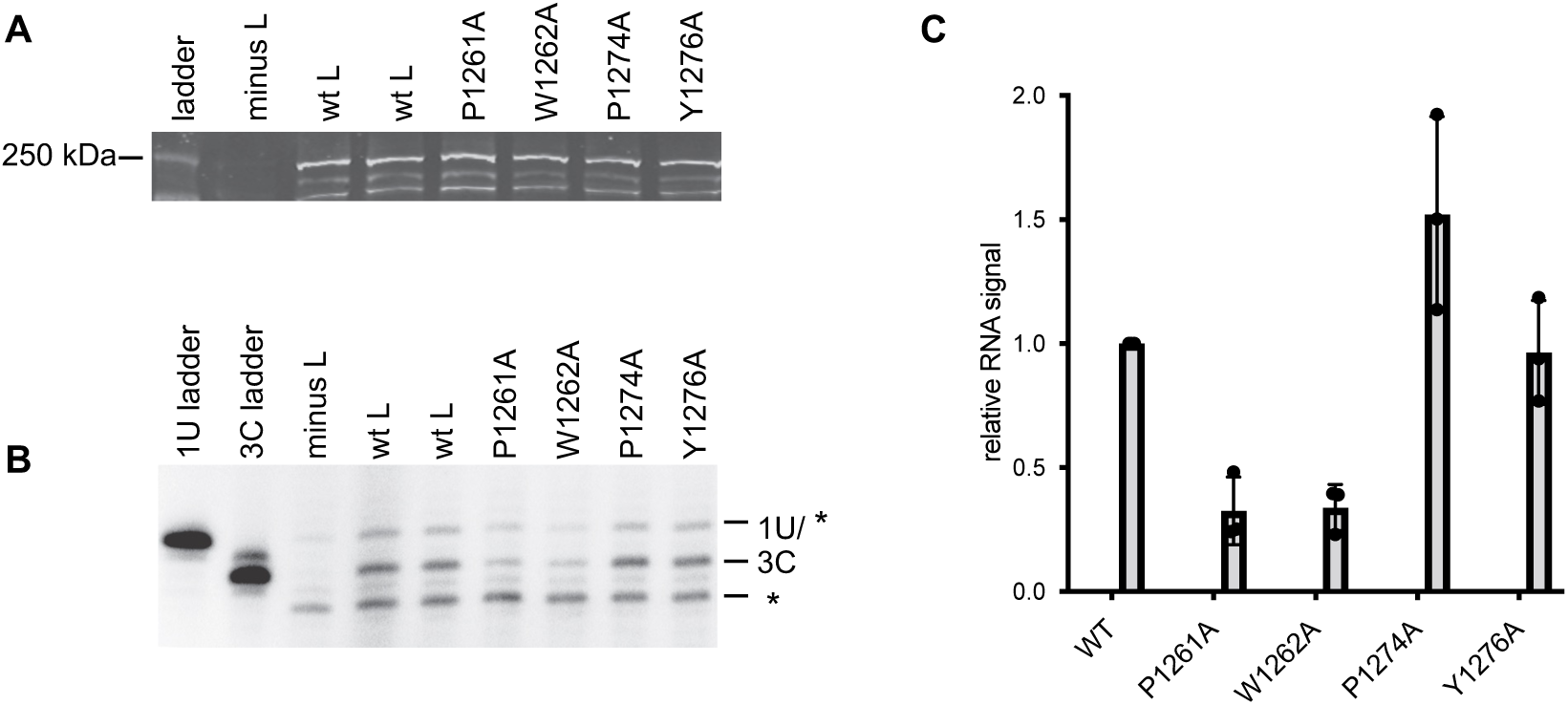
Analysis of L mutant protein activity at 30°C. (A) L mutant proteins containing an N-terminal FLAG tag were expressed in mammalian cells at 30°C. At 48 h post transfection, cell lysates were analyzed by Western blot analysis using a FLAG-specific antibody. The image is representative of three independent experiments. (B) Analysis of RNA products generated from the 1U and 3C sites in a minigenome containing a *le* promoter (shown in Fig 2A) by mutant L polymerases, using the same transfection conditions as used in Fig 2, except that the cells were incubated at 30°C rather than 37°C. The asterisks indicate non-specific background bands. (C) Quantification of the 3C initiation products presented in panel B. The bars show the mean and standard deviation for three independent experiments (the data points for each experiment are shown). Unfortunately, a background band that migrated at the same position as the 1U product prevented accurate quantification of this product. However, quantification of the 3C product showed that L mutants P1274A and Y1276A yielded RNA at a similar level as wt L, whereas the P1261A and W1262A mutants yielded RNA from the 3C initiation site at approximately 35% of wt levels. These results are similar to those obtained at 37°C (Fig 2).

